# Neuronal aging potentiates beta-amyloid generation via amyloid precursor protein endocytosis

**DOI:** 10.1101/616540

**Authors:** Tatiana Burrinha, Ricardo Gomes, Ana Paula Terrasso, Cláudia Guimas Almeida

## Abstract

Aging increases the risk of Alzheimer’s disease (AD). During normal aging synapses decline and β-Amyloid (Aβ) accumulates. An Aβ defective clearance with aging is postulated as responsible for Aβ accumulation, although a role for increased Aβ production with aging can also lead to Aβ accumulation. To test this hypothesis, we established a long-term culture of primary mouse neurons that mimics neuronal aging (lysosomal lipofuscin accumulation and synapse decline). Intracellular endogenous Aβ42 accumulated in aged neurites due to increased amyloid-precursor protein (APP) processing. We show that APP processing is up-regulated by a specific age-dependent increase in APP endocytosis. Endocytosed APP accumulated in early endosomes that, in turn were found augmented in aged neurites. APP processing and early endosomes up-regulation was recapitulated in vivo. Finally, we found that inhibition of Aβ production reduced the decline in synapses in aged neurons. We propose that potentiation of APP endocytosis by neuronal aging increases Aβ production, which contributes to aging-dependent decline in synapses.

**Summary:** How aging increases the risk of Alzheimer’s disease is not clear. We show that normal neuronal aging increases the intracellular production of β-amyloid, due to an upregulation of the amyloid precursor protein endocytosis. Importantly, increased Aβ production contributes to the aging-dependent synapse loss.

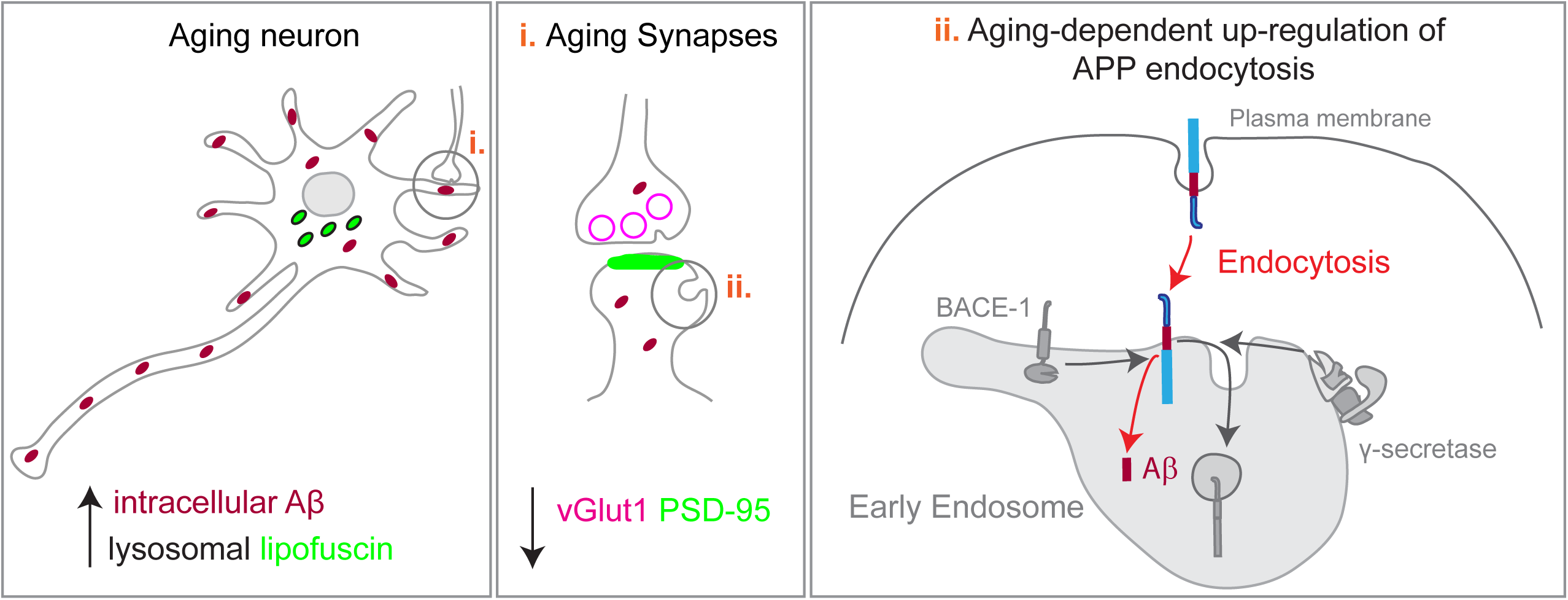

## Introduction

The prevalence of neurodegenerative diseases in the aging population is increasing due to the higher life expectancy. Alzheimer’s disease (AD) is the most common neurodegenerative disease and it remains without effective treatment. Cognitive decline develops with aging and often precedes AD (Yankner et al., 2008). The multifactorial mechanisms underlying this aging-associated cognitive decline are likely silent pathological mechanisms that will eventually trigger the onset of AD(Gauthier et al., 2006; Veitch et al., 2018). One of the mechanisms is the cellular aging of neurons, the main cellular target of AD (Mattson and Magnus, 2006). Unlike other brain cells, most neurons are born embryonically, do not undergo cell division, and thus have the organism chronological age. Neurons are the major producers of beta-amyloid (Aβ), the synaptotoxic agent that accumulates in amyloid plaques and triggers neurofibrillary tau tangle formation, the pathological hallmarks of AD(Cole and Vassar, 2007; Palop and Mucke, 2010; Selkoe and Hardy, 2016). Thus, neurons are not only targets of AD but also play a central role in the disease. For a long time, it was thought that, during aging, neurons would progressively die, underlying the cognitive decline that would lead to AD. However, it recently became apparent that initial cognitive decline results from synaptic dysfunction (Morrison and Baxter, 2012). Aging-synaptic decline has been characterized morphologically, by the loss of synapses and disrupted remodeling of spines, the postsynaptic compartments; and physiologically, by impaired synaptic plasticity, such as defects in potentiation and weakening of synapses, in hippocampus and prefrontal cortex, likely undermining establishment of new memories (Morrison and Baxter, 2012; Dickstein et al., 2013; Yankner et al., 2008; Samson and Barnes, 2013). How aging of neurons drives the loss of synapses associated with aging is unclear.

Identifying the causal mechanisms of aging-synaptic decline is urgent to devise treatments to reverse it, thus prevent cognitive decline and AD.

Neurons without the diluting effect of cell division accumulate damage, such as lipofuscin, an auto-fluorescent undegradable long-lived complex that consists of oxidatively damaged proteins and lipids, considered as the cellular aging hallmark (Mattson and Magnus, 2006). Interestingly, lipofuscin accumulates in lysosomes, the endpoint of endosomal trafficking pathway (Mattson and Magnus, 2006; Brunk and Terman, 2002).

From work done on familial AD, it has been established that the cascade that leads to AD is initiated by the progressive accumulation of Aβ. Aβ accumulation results from the imbalance between production and clearance. Aβ accumulation in familial AD is mostly due to increased Aβ production by neurons, caused by mutations that potentiate the cleavage of amyloid precursor protein (APP) by β-secretase (BACE1), or alter the cleavage by γ-secretase to produce more Aβ42. However, even in the absence of mutations, Aβ42 is normally produced (Haass et al., 1992) and it can accumulate in the brain with aging in mice, monkeys and humans (Baker-Nigh et al., 2015; Marks et al., 2017; Kikuchi et al., 2011; Lesné et al., 2013; Petersen et al., 2016; Blair et al., 2014). Importantly, the formation of extracellular amyloid plaques is preceded by a progressive intracellular accumulation of Aβ associated with synapse dysfunction (Gouras et al., 2005; Takahashi et al., 2002, 2004; Baker-Nigh et al., 2015; Welikovitch et al., 2018; Blair et al., 2014; Eimer and Vassar, 2013; Pensalfini et al., 2014; LaFerla et al., 2007).

It has been postulated that the clearance of Aβ is diminished with aging. Aβ clearance is likely decreased because neprilysin, one of the most important neuronal Aβ degrading enzymes, loses activity with aging (Iwata et al., 2002) and APOEe4, a variant in an Aβ binding lipoprotein associated with aging and AD, impedes an effective transport of Aβ out of the brain through the blood brain barrier (Castellano et al., 2011; Shibata et al., 2000). In contrast, the contribution of increased production to Aβ accumulation with aging is less clear. Evidence points to an aging-dependent alteration in BACE1 and γ-secretase activities (Guix et al., 2012; Fukumoto et al., 2004; Vassar et al., 2009), but not in APP expression (Flood et al., 1997; Gegelashvili et al., 1994). From studies of AD genetic risk factors, we and others have shown that alterations in the traffic of APP into endosomes enhances APP processing and the generation of Aβ (Andersen et al., 2006; Herskowitz et al., 2012; Rajendran and Annaert, 2012; Ubelmann et al., 2017b; Xiao et al., 2012). It is unknown if during aging the endocytic trafficking of APP is altered to increase Aβ production. Moreover, whether an aging-dependent Aβ accumulation, independent of Alzheimer mutations, impacts the maintenance of synapses, is unknown. In this study we have investigated whether changes in APP endosomal trafficking with neuronal aging contribute to increased Aβ production and whether the increased generation of Aβ is responsible for the age-dependent synaptic decline.

We chose primary mouse post-mitotic cortical neurons aged in culture as a model of neuronal aging since these cultures undergo *in vitro* an accelerated stereotyped process of differentiation and in four weeks recapitulate important aspects of *in vivo* aging such as lipofuscin accumulation, increased reactive oxygen species, protein oxidation and lipid alterations (Aksenova et al., 1999; Goslin and Banker, 1989; Martin et al., 2011; Youmans et al., 2012; Papa et al., 1995; Trovò et al., 2013). We found that aged primary neurons accumulate intracellular Aβ42 which correlates with the increase in APP processing. Mechanistically, such phenomenon could be due to our discovery that APP endocytosis is up-regulated in aged neurons, especially in neurites. We validated the up-regulation of APP processing and endocytosis in aged brain. Finally, we established causality between Aβ production and synapse decline since we observed that blocking APP processing increased the number of synapses in aged neurons. Overall, we identified a mechanism whereby neuronal aging may be contributing to the development of Alzheimer’s disease.

## Results

### Intracellular Aβ42 production increases with neuronal aging

Primary embryonic cortical neurons in culture undergo a stereotyped differentiation process. Briefly, during the first 7 days *in vitro* (7 DIV), axons and dendrites are formed; in the second week (14 DIV) dendrites develop completing maturation after three weeks (21 DIV) (Goslin and Banker, 1989; Boyer et al., 1998). In accordance, we observed the maximum expression of microtubule associated protein (MAP2), a marker of neuronal differentiation, at 21 DIV (Fig. 1A, B). After one more week in culture, several groups have reported that neurons start aging, or cellular senescence (Aksenova et al., 1999; Goslin and Banker, 1989; Martin et al., 2011; Youmans et al., 2012; Papa et al., 1995; Trovò et al., 2013). We observed that, primary neurons cultured for 28 DIV had levels of MAP2 similar to 21 DIV neurons (Fig. 1A, B) and did not present gross morphological changes such as axonal bead-like degeneration or dendritic shrinkage (Fig.1C-F). Importantly, 28 DIV neurons showed canonical signs of cellular aging, such as the formation of auto-fluorescent aging granules, or lipofuscin (Fig. S1A-E) as previously described (Gray and Woulfe, 2005). Importantly, the number of auto-fluorescent granules within lysosome associated membrane protein 1 (LAMP1)-positive lysosomes was significantly higher in the cell bodies of aged neurons (28 DIV) than in mature neurons (21 DIV), consistent with the aging-dependent accumulation of lysosomal lipofuscin (Fig. 1G-K). Thus, we established a model of cellular aging of post-mitotic neurons that, after reaching maturation, undergo cellular aging, without neurodegeneration.

**Figure 1.**
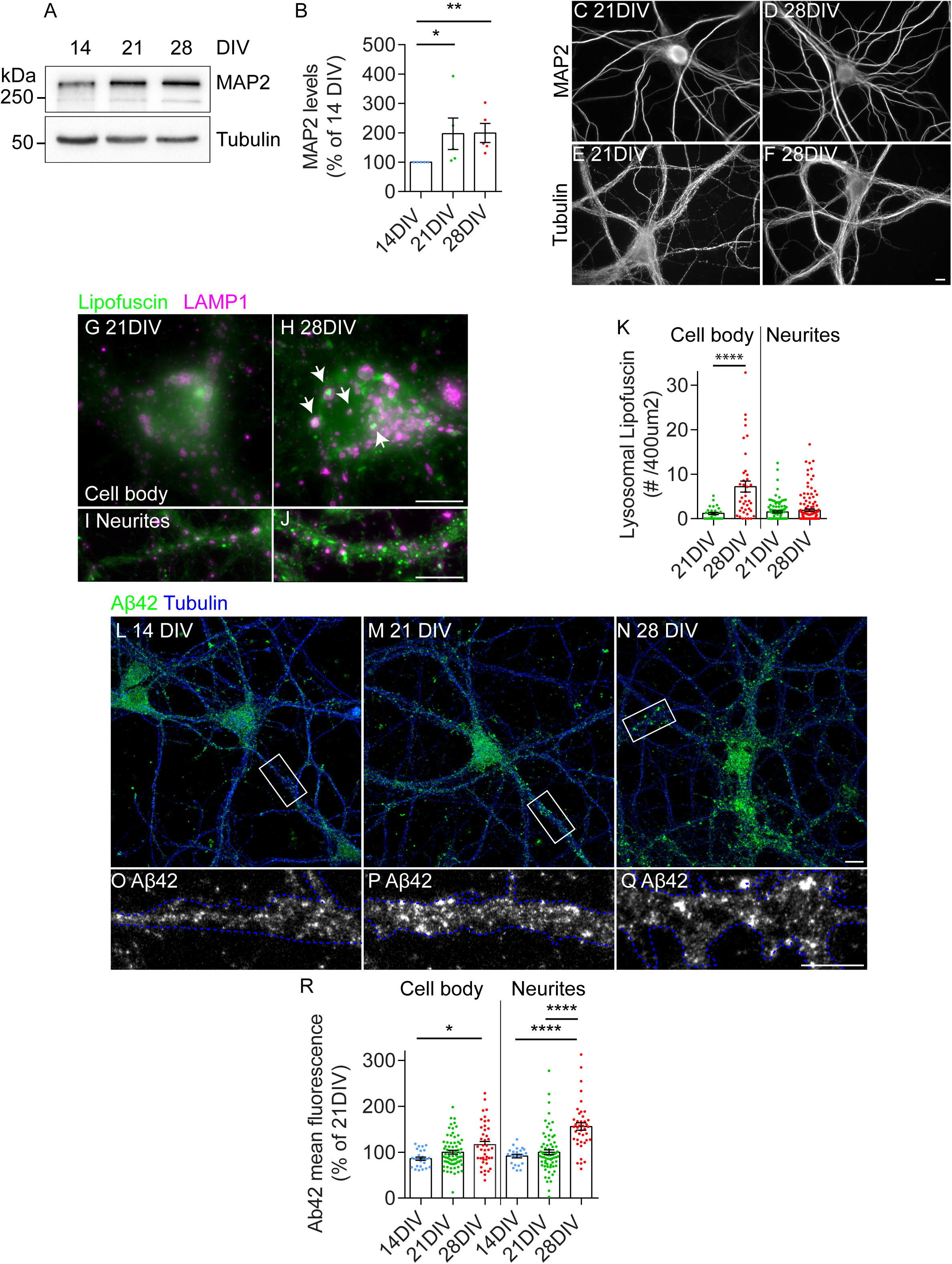
Primary neurons aged in culture accumulate Aβ42. A. MAP2 total levels by western-blot of wild-type mouse primary cortical neurons after 14 days in vitro (DIV), 21 DIV and 28 DIV. Tubulin was immunoblotted as loading control. B. Quantification of MAP2 levels normalized to tubulin and to percentage of 14 DIV neurons (n= 5; *P = 0.0432 21 DIV vs. 14 DIV neurons, **P = 0.0095 28 DIV vs. 14 DIV neurons, one-way ANOVA on ranks with *post hoc* Dunn’s testing, mean ± SEM). C-F. Representative images of the morphology of neurons after 21 DIV and 28 DIV, immunolabelled with anti-MAP2 (C, D) and anti-tubulin (E, F), analyzed by epifluorescence microscopy. Scale bars, 10 µm. G-J. Lipofuscin (arrowheads; green) and LAMP1 (magenta) localization in cell bodies (G, H) and neurites (I, J) of 21 DIV and 28 DIV neurons labelled with anti-Lamp1 and analyzed by epifluorescence microscopy. Lipofuscin was identified by the presence of auto-fluorescent granules in Lamp1-positive lysosomes in cell bodies. Scale bars, 10 µm. K. Quantification of the number of lysosomal lipofuscin, defined by the colocalization of auto-fluorescent granules with Lamp1-positive lysosomes, per area (400 µm2) in cell bodies and neurites (n=4, N_cellbody_ = 28-39, N_neurites_= 104-129, ****P_cellbody_ < 0.0001 28 DIV vs. 21 DIV, Mann-Whitney test, mean ± SEM). L-Q. Intracellular endogenous Aβ42 (green) and tubulin (blue) in 14 DIV, 21 DIV and 28 DIV, immunolabelled with anti-Aβ42 (clone 12F4) and anti-tubulin, analyzed by confocal microscopy. The white rectangles indicate the neurites magnified below (O-Q). Outlined neurites based on tubulin (blue) showing Aβ42 punctate staining. Scale bars, 10 µm. R. Quantification of Aβ42 (12F4) intensity in cell body and neurites (n = 3-6, N_cellbody_ = 25-70, N_neurites_= 22-73, *P_cellbody_ = 0.0158 28 DIV vs. 14 DIV, ****P_neurites_ < 0.0001 28 DIV vs. 14 DIV, ****P_neurites_ < 0.0001 28 DIV vs. 21 DIV, one-way ANOVA on ranks with *post hoc* Dunn’s testing, mean ± SEM).

Previously, we demonstrated that intracellular Aβ42 progressively accumulates in older primary cortical neurons due to overexpression of mutant hAPP (Swe) (Takahashi et al., 2004). Now, to determine if neuronal aging can induce quantitative changes in intracellular endogenous Aβ42, we used a novel and highly sensitive semiquantitative immunofluorescence assay using an anti-Aβ42 C-terminal specific antibody (12F4) (Ubelmann et al., 2017b). By analyzing the accumulation of Aβ42 in neurites and cell bodies separately we discovered a significant increase in Aβ42 accumulation (56%) in aged neurites compared to mature neurites (Fig. 1M, N, P, Q, R). In the aged cell bodies, Aβ42 increased (30 %) less than in aged neurites, being significantly different only when compared with immature neurites (14 DIV) (Fig. 1L, O, R). These results indicate that neuronal aging drives intracellular Aβ42 accumulation specifically in neurites.

### APP processing is increased with aging

To assess if neuronal aging increased Aβ42 by potentiating the processing of APP we analyzed the levels of APP C-terminal fragments (APP CTFs) by western blot using an anti-APP C-terminal (APPY188) antibody. We found that the levels of APP CTFs increased in aged neurons compared with those in mature neurons (Fig. 2A, B). This increase in APP CTFs together with the increase in intracellular Aβ42 indicate that endogenous APP is being more processed in aged neurons. Of note, we found that neuronal maturation did not significantly affect APP processing, since the levels of APP CTFs in mature neurons (21 DIV) were similar to those in immature neurons (14DIV) (Fig. 2A, B). Interestingly, the levels of full-length APP were not significantly altered between mature and aged neurons (Fig. 2C, D), discarding a major impact of neuronal aging in culture on APP expression or degradation. We observed an increase in the levels of APP when comparing mature to immature neurons, likely due to APP being increasingly expressed with neuronal maturation and synaptogenesis (Fig. 2C, D) (Hung et al., 1992; Nicolas and Hassan, 2014). Moreover, we found the levels of BACE1 and nicastrin, a γ-secretase subunit, unaltered in aged neurons when compared with mature neurons (Fig. 2E, F), recapitulating *in vivo* data (Fukumoto et al., 2004; Guix et al., 2012). The increase in APP processing without changes in the full-length APP indicate that APP trafficking may be altered with neuronal aging.

**Figure 2.**
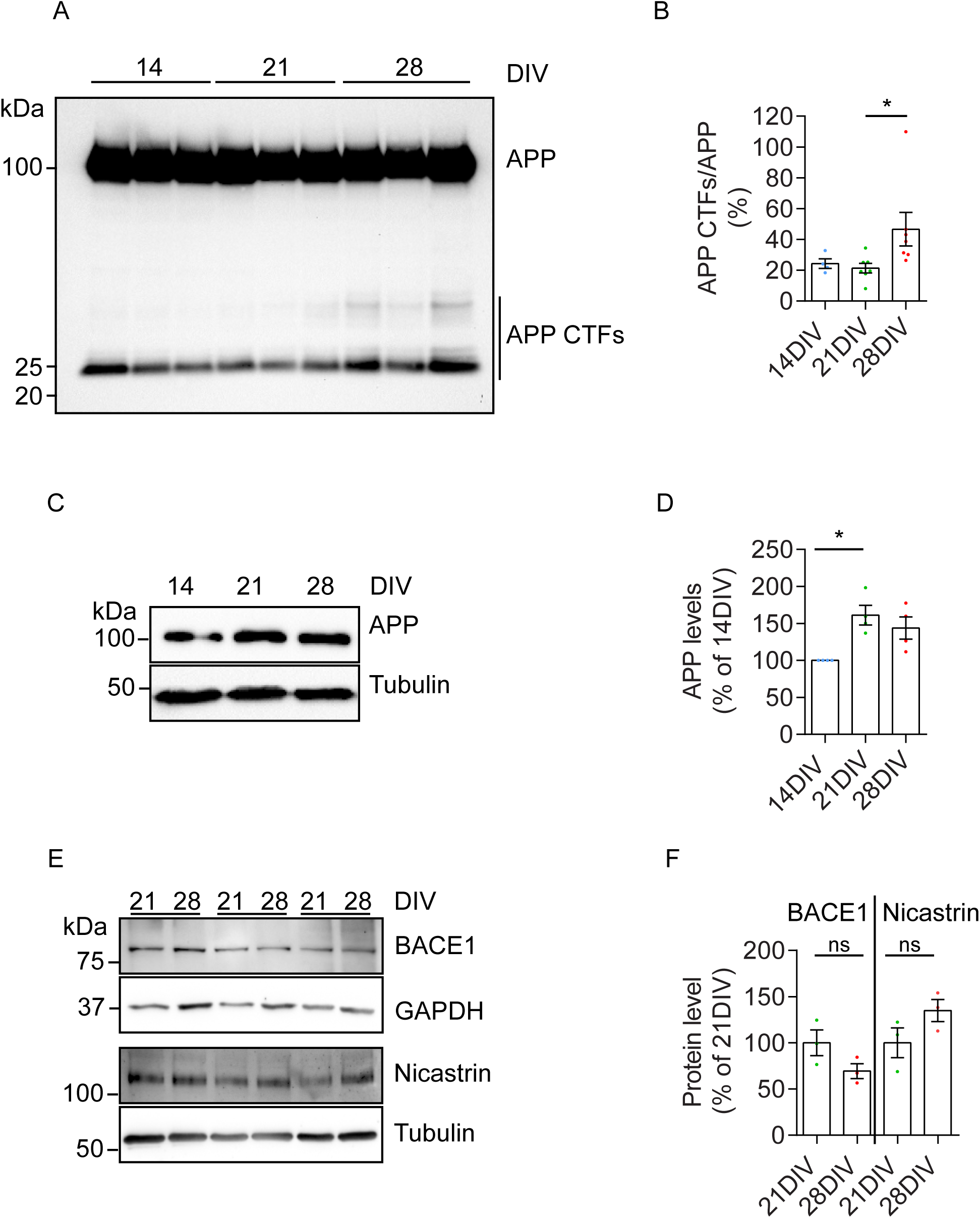
Neuronal aging potentiates the processing of APP. A. Endogenous APP and APP-CTFs levels in neurons at 14 DIV, 21 DIV and 28 DIV, analyzed by western blot with anti-APP antibody (Y188). B. Quantification of APP-CTFs levels normalized to APP (n = 4-7; \**P* = 0.0152 28 DIV vs. 21 DIV neurons, one-way ANOVA on ranks with post hoc Dunn’s testing, mean ± SEM). C. Endogenous APP in neurons at 14 DIV, 21 DIV and 28 DIV analyzed by Western blot with anti-APP antibody (Y188). Tubulin was immunoblotted as loading control. D. Quantification of APP levels normalized to percentage of 14 DIV neurons (n = 4; \**P* = 0.0284 21 DIV vs. 14 DIV neurons, one-way ANOVA on ranks with post hoc Dunn’s testing, mean ± SEM). E. Endogenous BACE1 analyzed by Western blot with anti-BACE1 antibody and GAPDH as loading control of neurons at 21 DIV and 28 DIV (top panels). Endogenous subunit of the gamma-secretase complex, nicastrin, analyzed by Western blot with anti-nicastrin antibody and tubulin as loading control of neurons at 21 DIV and 28 DIV (bottom panels). F. Quantification of BACE1 and nicastrin levels normalized to percentage of 21 DIV neurons (n = 3; 21 DIV were not significant different from 28 DIV (ns), Wilcoxon test, mean ± SEM).

### Intracellular APP is enriched in neurites of old neurons

Since Aβ accumulation and APP processing occur mostly in neurites, close to distal synapses (Takahashi et al., 2004; Das et al., 2016) we analyzed the impact of neuronal aging on the distribution of APP between the cell body and neurites. We applied our semiquantitative immunofluorescence assay to measure the enrichment of APP in neurites vs cell bodies of aged and mature neurons. We found a major increase in APP intensity in aged neurites (52%) compared to mature neurites (Fig. 3A-C). In contrast, no significant difference in APP intensity was observed with neuronal aging in the cell body (Fig. 3A-C). The increase of APP in neurites correlated with the increased level of Aβ42 detected in neurites (Fig. 1N, Q, R). Next, we explored if this increase was occurring in axons or dendrites. We measured APP in axons identified by Ankyrin G (AnkG positive) and in dendrites (AnkG negative) (Fig. 3D, E). We found that APP is polarized to axons in both mature and aged neurons (Fig. 3F-3I, 3K). Quantification of the ratio of APP intensity in axons over dendrites showed that the axonal polarization of APP is slightly but significantly decreased (15%) in aged neurons compared to mature neurons, indicating that APP distributed more to dendrites than to axons of aged neurons (Fig. 3J). Indeed, we found that APP increased more in dendrites (42 %) than in axons (28 %) with neuronal aging (Fig. 3K).

**Figure 3.**
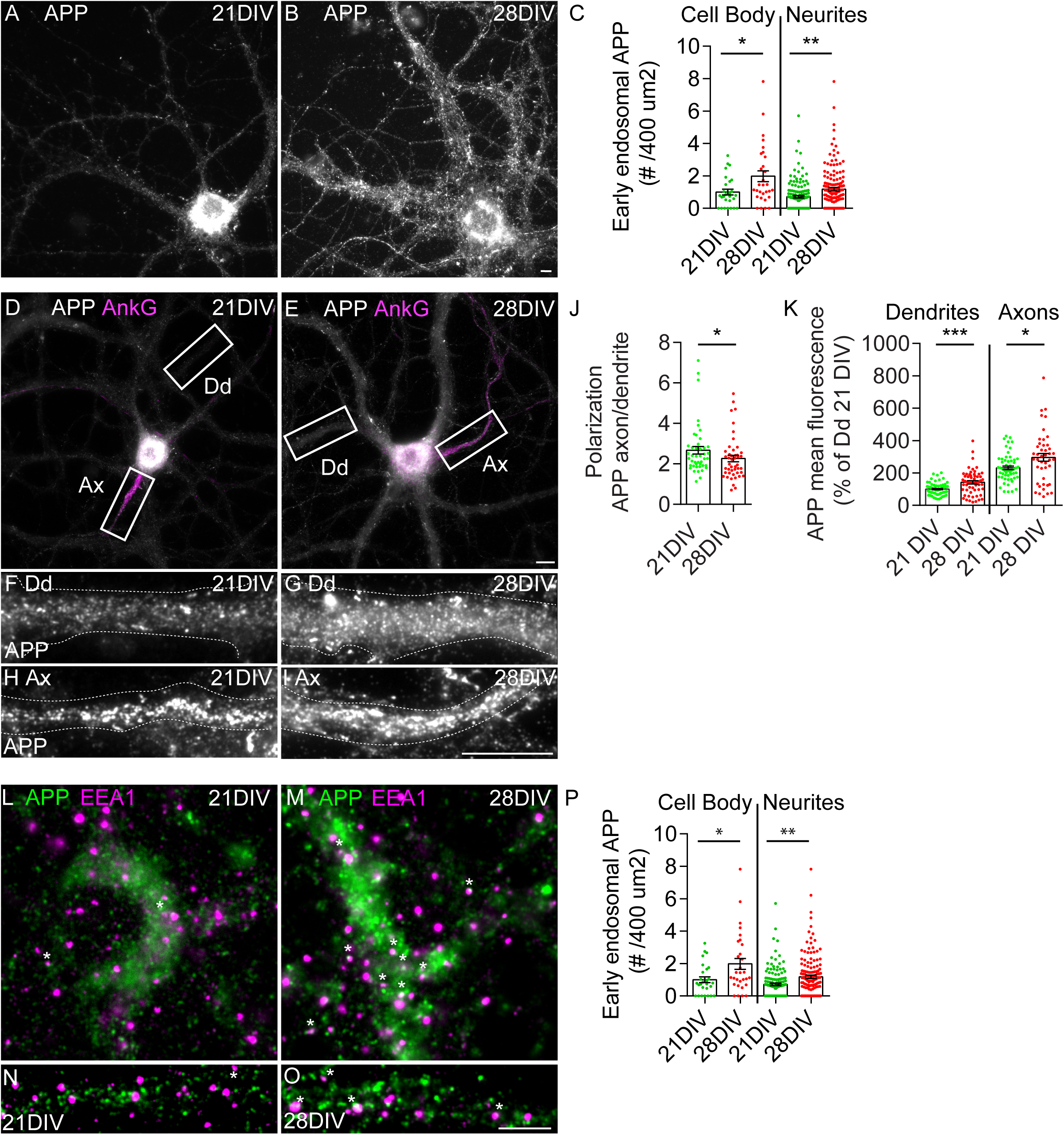
Neuronal aging polarizes APP distribution towards neurites. A-C. APP localization in neurons detected by immunofluorescence with anti-APP antibody (Y188) of neurons at 21 DIV (A) and 28 DIV (B), analyzed by epifluorescence microscopy. Scale bar, 10 µm. (C) Quantification of the mean intensity of APP using ImageJ “measure” function in cell bodies and neurites of 21 and 28 DIV neurons. Results were normalized to percentage of 21 DIV. (n = 3, N_cellbody_=28-30; N_neurites_= 29-30; \*\*\*\**P* < 0.0001 28 DIV vs. 21 DIV neurites, unpaired t-test, mean ± SEM). D-K. APP localization in axons and dendrites by immunofluorescence at 21 DIV (D) and 28 DIV (E), with anti-APP antibody (Y188) and anti-ankyrinG (AnkG; magenta) to identify axons, analyzed by epifluorescence microscopy. The white rectangles indicate the magnified dendrites (F and G) and axons (H and I). Scale bar, 10 µm. (J) Quantification of the axon/dendrite ratio of APP calculated to quantify APP polarization is shown (n=3, N_21DIV_ = 44, N_28DIV_ = 46; \**P* = 0.0322, Mann-Whitney test; mean ± SEM). (K) Quantification of the mean intensity of APP using ImageJ “measure” function in dendrites and axons of 21 and 28 DIV neurons is shown. Results were normalized to percentage of 21 DIV dendrites. (n = 4-5, N_dd_=57-76; N_axon_= 44-55; \*\*\**P* = 0.0002 28 DIV vs. 21 DIV dendrites, \**P* = 0.0258 28 DIV vs. 21 DIV axons, Mann-Whitney test, mean ± SEM). L-P APP localization in EEA1-positive early endosomes in the cell body (L and M) and in neurites (N and O) by immunofluorescence of neurons at 21 DIV and 28 DIV, with anti-APP antibody (Y188; green) and anti-EEA1 (EEA1; magenta), analyzed by epifluorescence microscopy and the images are displayed after background subtraction with Fiji. Scale bar, 10 µm. (P) Quantification of the number of APP positive early endosomes at steady state per 400 µm2 of cell body or neurite of 21 DIV and 28 DIV neurons using ICY “Colocalizer” protocol (n=3; N_cellbody_=28-30; N_neurites_= 131-144; \**P =* 0.0264 28 DIV vs. 21 DIV cell bodies, \*\**P =* 0.0012 28 DIV vs 21 DIV neurites, Mann Whitney test, mean ± SEM).

Since early endosomes are the main site for APP processing in neurons, we investigated if APP is more localized to early endosomes in aged neurons. To do so, we analyzed APP colocalization with EEA1, an early endosome marker, in cell bodies (Fig. 3L, M) and neurites (Fig. 3N, O) of aged and mature neurons. We automatically measured the number of endosomes containing APP and found more APP in early endosomes in cell bodies and neurites of aged neurons compared to mature neurons (Fig. 3L-P). These results indicate that, despite the overall unaltered levels of APP (Fig. 2C, D) there is an increase in the APP levels at steady state in neurites (Fig. 3C) and in early endosomes (Fig. 3P).

### APP endocytosis increases with aging

To further explore the age-dependent increase of APP in early endosomes we went on to directly assess APP endocytosis. Since APP endocytosis is required for Aβ production, we investigated if it is potentiated by neuronal aging. To study APP endocytosis, we used two endocytosis assays. First, we performed a bulk surface proteins internalization assay (Fig. 4A). In this assay, we biotinylated live neurons with cleavable NHS-SS biotin. Biotinylated APP was detected using a specific anti-APP antibody (Y188)(Ubelmann et al., 2017b). The cleavage of surface biotin with non-permeable reducing agent glutathione (GSH) allowed for the specific detection of endocytosed biotinylated APP (Snyder et al., 2005). After 10 min chase, the majority of biotinylated APP remained at the cell surface (10’-GSH) and only a small fraction of surface APP was endocytosed (10’ + GSH). After 30 min chase, the level of endocytosed APP (30’ + GSH) increased, being more endocytosed by aged neurons than by mature neurons (Fig. 4B). Quantification revealed that, APP endocytosis at 10 min was 7-fold higher in aged neurons than in mature neurons. After 30 min, aged neurons APP endocytosis was even higher (10-fold) (Fig. 4C). This increase in endocytosis was not due to an increase in surface APP in aged neurons since it was not significantly different from that in mature neurons (Fig. 4D), neither did it translate into higher degradation of APP since APP total levels in aged neurons were not significantly different from that in mature neurons (Fig. 4E, Fig. 2D).

**Figure 4.**
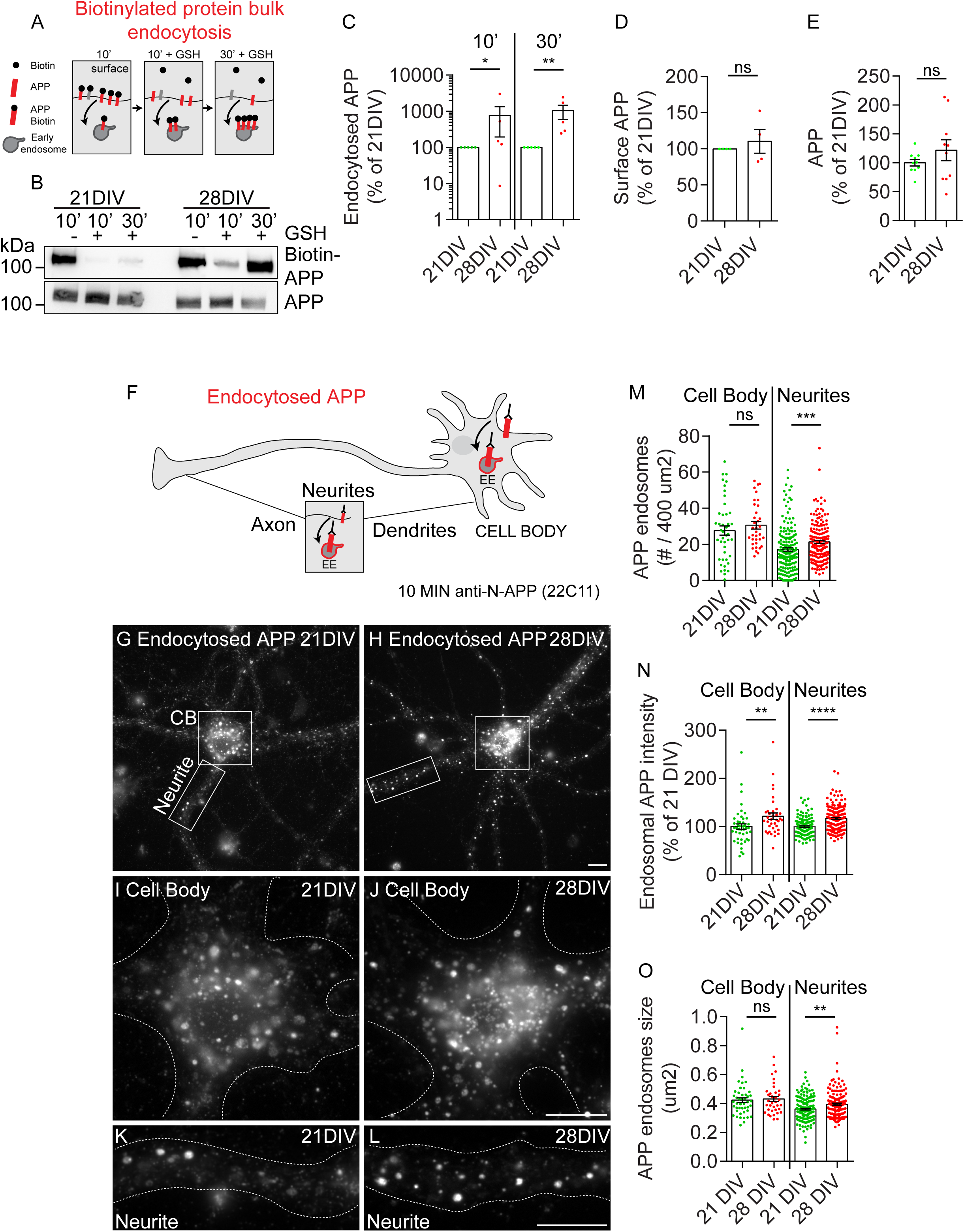
Aged neurons endocytosed more APP. A. Scheme illustrating the endocytosis assay using biotinylation of surface proteins. Biotinylated proteins after 10 min chase (10’); Endocytosed biotinylated proteins after 10 min chase and removal of surface biotin with non-cell permeable glutathione (GSH) (10’+ GSH); Endocytosed biotinylated proteins after 30 min chase and removal of surface biotin with non-cell permeable glutathione (GSH) (30’+GSH). B. Surface and endocytosed biotin-APP (see A.) in neurons at 21 DIV and 28 DIV. Biotinylated (Biotin-APP) and total APP were detected with anti-APP (Y188) by western blot. C. Quantification of endocytosed biotin-APP normalized to percentage of 21 DIV, results are shown in logarithmic scale (n = 5, \**P*_10’_ = 0.0476 28 DIV vs 21 DIV, \*\**P*_30’_ = 0.0079 28 DIV vs 21 DIV, Kolmogorov-Smirnov test; mean ± SEM). D. Quantification of surface biotin-APP normalized to percentage of 21 DIV, results are shown in logarithmic scale (n = 4, Not significant *P* = 0.2500 28 DIV vs 21 DIV, Wilcoxon test; mean ± SEM). E. Quantification of total APP normalized to percentage of 21 DIV, results are shown in logarithmic scale (n = 5, not significant *P* = 0.4316 28 DIV vs 21 DIV, Wilcoxon test; mean ± SEM). F-L. Scheme illustrating the APP endocytosis assay using an antibody specific against the N-terminus of APP (22C11). During a 10 min pulse the antibody binds surface APP and internalizes into early endosomes. APP endocytosis in neurons at 21 DIV (G) and 28 DIV (H). White squares indicate the magnified cell bodies shown in (I) and (J). The white rectangles indicate the magnified neurites shown in (K) and (L). Scale bars, 10 µm. M. Quantification of the number of APP endosomes, defined by the 10 min uptake of 22C11, per area (400 µm2) in cell bodies and neurites (n=5, N_cellbody_ = 34-43, N_neurites_= 167-175, not significant, *P*_cellbody_ = 0.3520 28 DIV vs. 21 DIV, \*\*\**P*_neurites_ = 0.0003 28 DIV vs. 21 DIV, Mann-Whitney test, mean ± SEM). N. Quantification of the mean intensity of APP endosomes using Icy “spot detector” plugin in cell body and neurites of 21 and 28 DIV neurons is shown. Results were normalized to percentage of 21 DIV (n =5, N_cellbody_= 36-45; N_neurites_=167-170; \*\**P*_cellbody_ = 0.0044 28 DIV vs. 21 DIV, \*\*\*\**P*_neurites_ = 0.0001 28 DIV vs. 21 DIV Mann-Whitney test, mean ± SEM). Q. Quantification of the mean size of each APP endosomes (µm2) using Icy “spot detector” plugin in cell body and neurites of 21 and 28 DIV neurons (n =5, N_cellbody_= 36-45; N_neurites_=161-167; not significant *P*_cellbody_ = 0.7071 28 DIV vs. 21 DIV, \*\**P*_neurites_ = 0.0026 28 DIV vs. 21 DIV neurites, Mann-Whitney test, mean ± SEM).

Next, since we observed a redistribution of APP to neurites, we wanted to investigate if APP endocytosis was increased in neurites of aged neurons. For that, we used an antibody internalization assay by pulsing live neurons with a mouse monoclonal antibody against the extracellular N-terminal domain of APP (22C11). After 10 min, we detected endocytosed APP with a fluorescently labelled secondary anti-mouse antibody to detect the endocytosed 22C11 bound to APP (Ubelmann et al., 2017a)(Fig. 4F). Of note, we were surprised by the prompt detection of 22C11 endocytosis in mature and aged neurons since in previous work using immature neurons (9 DIV), APP overexpression was required to consistently detect APP endocytosis (Ubelmann et al., 2017b). This indicates that the levels of surface APP, or the levels of APP endocytosis possibly increase with neuronal maturation allowing us to follow the endocytosis of endogenous APP. We colocalized endocytosed APP with Rab5, a Rab GTPase specific of early endosomes, confirming the localization of endocytosed APP to early endosomes (Fig. S2). We observed that, in mature neurons, APP endosomes were more easily detected in the cell bodies than in neurites, while in aged neurons APP endosomes were also easily detected in neurites (Fig. 4G-4L). To quantify APP endocytosis, we segmented each APP endosome and analyzed its density, intensity and size in cell bodies and neurites. We found that the density, intensity and size of APP endosomes increased in aged neurites when compared to mature neurites (Fig. 4M-4O). In contrast, only the intensity of endocytosed APP increased in the cell bodies of aged neurons when compared to mature neurons (Fig. 4N). The number of APP positive endosomes was larger than observed at steady state (Fig. 3P) likely reflecting that only a small fraction of the total cellular APP undergoes endocytosis. Together these results indicate that APP endocytosis increases with aging mainly in neurites thus explaining the important rise observed in Aβ42 neuritic accumulation (Fig. 1Q, R).

### Neuronal aging leads to early endosomes increase

Early endosomes receive newly endocytosed cargo and are the major site of APP processing and Aβ production. Thus, we wondered if the increase in endocytosed APP was accompanied by alterations in early endosomes.

To analyze early endosomes, we immunolabelled neurons with anti-EEA1 antibody, and quantified their density, size and intensity in cell bodies and neurites of mature and aged neurons. We found that early endosomes were overall bigger and brighter in aged neurons and especially in neurites (Fig. 5A-5H), but their density was unaltered (Fig. 5I), indicating that early endosomes do not increase in number but are bigger by probably receiving more endocytosed APP with neuronal aging. This enlargement of early-endosomes was accompanied by an increase in EEA1 total levels in aged neurons as assessed by western-blot (Fig. 5K, L). The fact that the increased number of APP endosomes did not correspond to increased number of early endosomes could relate to APP being endocytosed into a subset of early endosomes, or that other cargo is accumulating in aged neurons.

**Figure 5.**
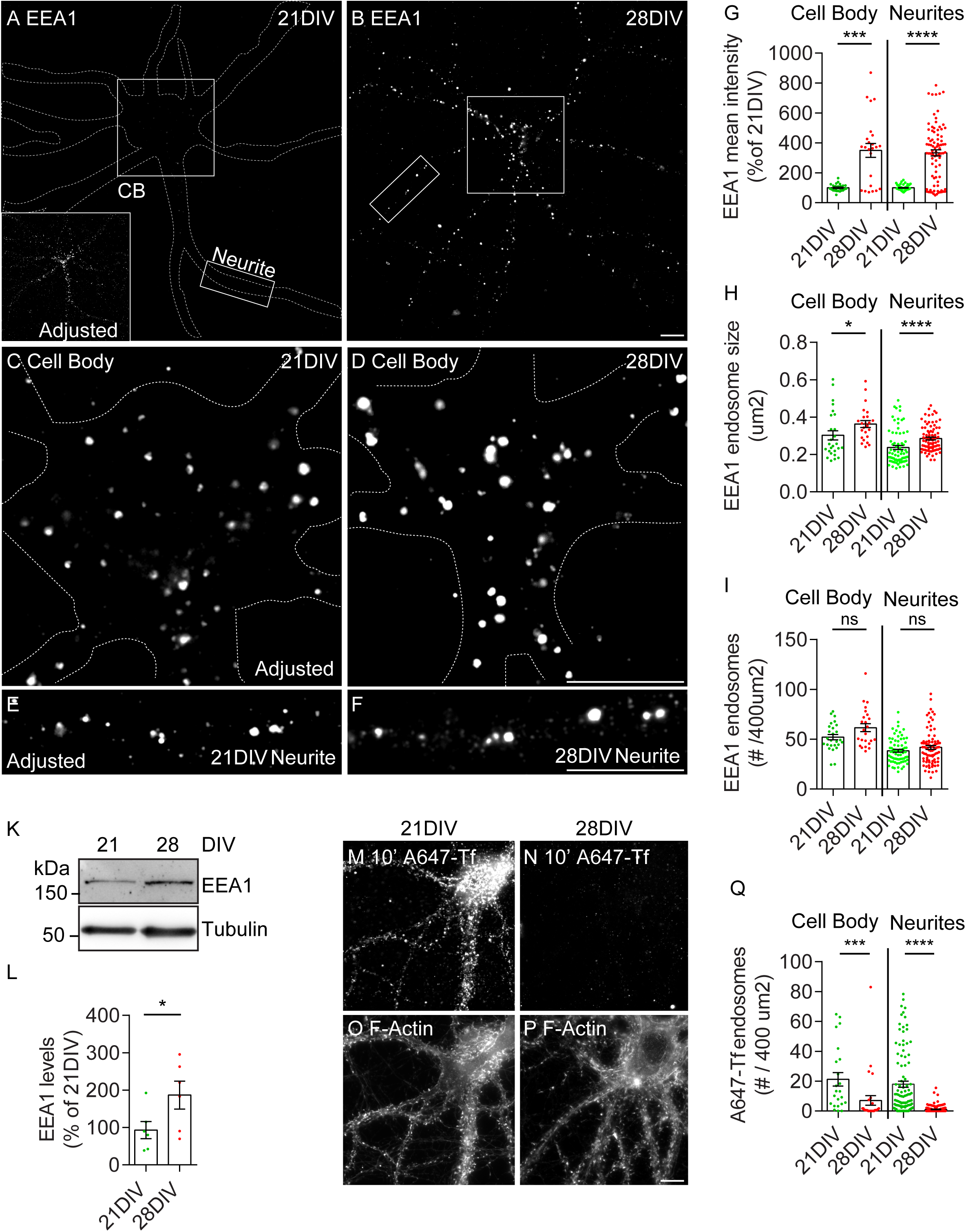
Aged neurons accumulate early endosomes. A-F. EEA1-positive endosomes detected by immunofluorescence with anti-EEA1 antibody of neurons at 21 DIV (A) and 28 DIV (B), analyzed by epifluorescence microscopy. Images are displayed after background subtraction with Fiji and neurons outlined in white. Inset shows EEA1-positive endosomes upon levels adjustment. White squares indicate the magnified cell bodies shown in (C) and (D). The white rectangles indicate the magnified neurites shown in (E) and (F). Note: in (C) and (E) the levels were adjusted to allow for comparison in the size and number of endosomes. Scale bars, 10 µm. G. Quantification of the mean intensity of each EEA1-positive early endosome, normalized to percentage of 21 DIV, in cell bodies and neurites of 21 DIV and 28 DIV neurons (n=3, N_cellbody_ = 24-26, N_neurites_= 72-85, ***P_cellbody_ = 0.0005 28 DIV vs. 21 DIV neurons, ****P_neurites_ < 0.0001 28 DIV vs. 21 DIV neurons, Mann-Whitney test, mean ± SEM). H. Quantification of the mean size of each EEA1-positive early endosome (µm2) in cell bodies and neurites of 21 DIV and 28 DIV neurons (n=3, N_cellbody_ = 24-26, N_neurites_= 72-85, *P_cellbody_ = 0.0131 28 DIV vs. 21 DIV, ****P_neurites_ < 0.0001 28 DIV vs. 21 DIV, Mann-Whitney test, mean ± SEM). I. Quantification of number EEA1-positive early endosomes per 400 µm2 of cell bodies and neurites of 21 DIV and 28 DIV neurons (n=3, N_cellbody_ = 24-26, N_neurites_= 72-85, ***P_cellbody_ = 0.0005 28 DIV vs. 21 DIV, ****P_neurites_ < 0.0001 28 DIV vs. 21 DIV, Mann-Whitney test, mean ± SEM). K. EEA1 levels analyzed by Western blot with anti-EEA1 antibody and tubulin as loading control of neurons at 21 DIV and 28 DIV. L. Quantification of EEA1 levels normalized to tubulin and to percentage of 21 DIV neurons (n = 6; *P = 0.0313 28 DIV vs. 21 DIV, Wilcoxon test, mean ± SEM). M-P. Transferrin endocytosis in neurons pulsed with Alexa647-Tf for 10’ at 21 DIV (M) and 28 DIV (N) neurons. Phalloidin was used to label F-actin in 21 DIV (O) and 28 DIV (P) neurons. Neurons were analyzed by epifluorescence microscopy. Scale bars, 10 µm. Q. Quantification of the number of transferrin endosomes per 400 µm2 in cell bodies and neurites. (n = 3, N_cellbody_ = 23-28, N_neurites_= 105-156, ***P_cellbody_ = 0.0002 28 DIV vs. 21 DIV, ****P_neurites_ < 0.0001 28 DIV vs. 21 DIV, Mann-Whitney test, mean ± SEM).

To assess this last hypothesis, we measured the endocytosis of transferrin, a canonical endocytic cargo. Similarly, we pulsed live neurons for 10 min with fluorescently labelled transferrin (A647-Tf) that upon binding to surface transferrin receptor is internalized and delivered to early endosomes (Maxfield and McGraw, 2004). We observed that endocytosis of transferrin was reduced in aged neurons, both in cell body and in neurites (Fig. 5M-5Q). This finding is supported by a previously described reduced kinetics of transferrin internalization due to defective clathrin recycling (Blanpied et al., 2003), implicated in the formation of endocytic vesicles. In addition, we cannot exclude that the transferrin receptor could be reduced at the surface of aged neurons and thus less available to endocytose transferrin. In either case, this result indicates that the endocytic mechanism used by APP is different from that used by transferrin. These results indicate that there is not a general upregulation of endocytosis with neuronal aging but rather a specific aging-dependent up-regulation of APP endocytosis.

### APP processing and early endosomes are up-regulated with brain aging

Seeking *in vivo* confirmation for our main mechanistic findings, we analyzed APP processing and the level of EEA1 in the cortex of aged (18 months) and adult (6 months) wild-type mice (C57BL/6) by western-blot (Fig. 6A-D). We found that the levels of APP CTFs (< 15 kDa) increased in aged mice compared to young mice (Fig. 6A, B). The bottom APP CTF band likely corresponds to both the non-amyloidogenic α-CTF (C83; 10 kDa), product of alpha-secretase and the amyloidogenic β-CTF (C89; 12 kDa), product of beta-secretase cleavage. The top APP CTF band (*) observed likely correspond to the longer amyloidogeneic β-CTF (C99; 14kDa) in two of the aged mice and in one young mouse. In addition, a higher molecular weight CTF ($; >25kDa) increased in two aged mice, likely corresponding to η-CTF (Fig. 6A) (Willem et al., 2015). Densitometric analysis of the levels of all APP CTFs showed a significant increase, indicating that APP processing increases in aged mice (Fig. 6B). Importantly, we did not detect alterations in the levels of full-length APP *in vivo* (Fig. 6C). Again, as in *in vitro* aged neurons, we found that the levels of EEA1 increased in aged mice when compared to young mice (Fig. 6A, D).

**Figure 6.**
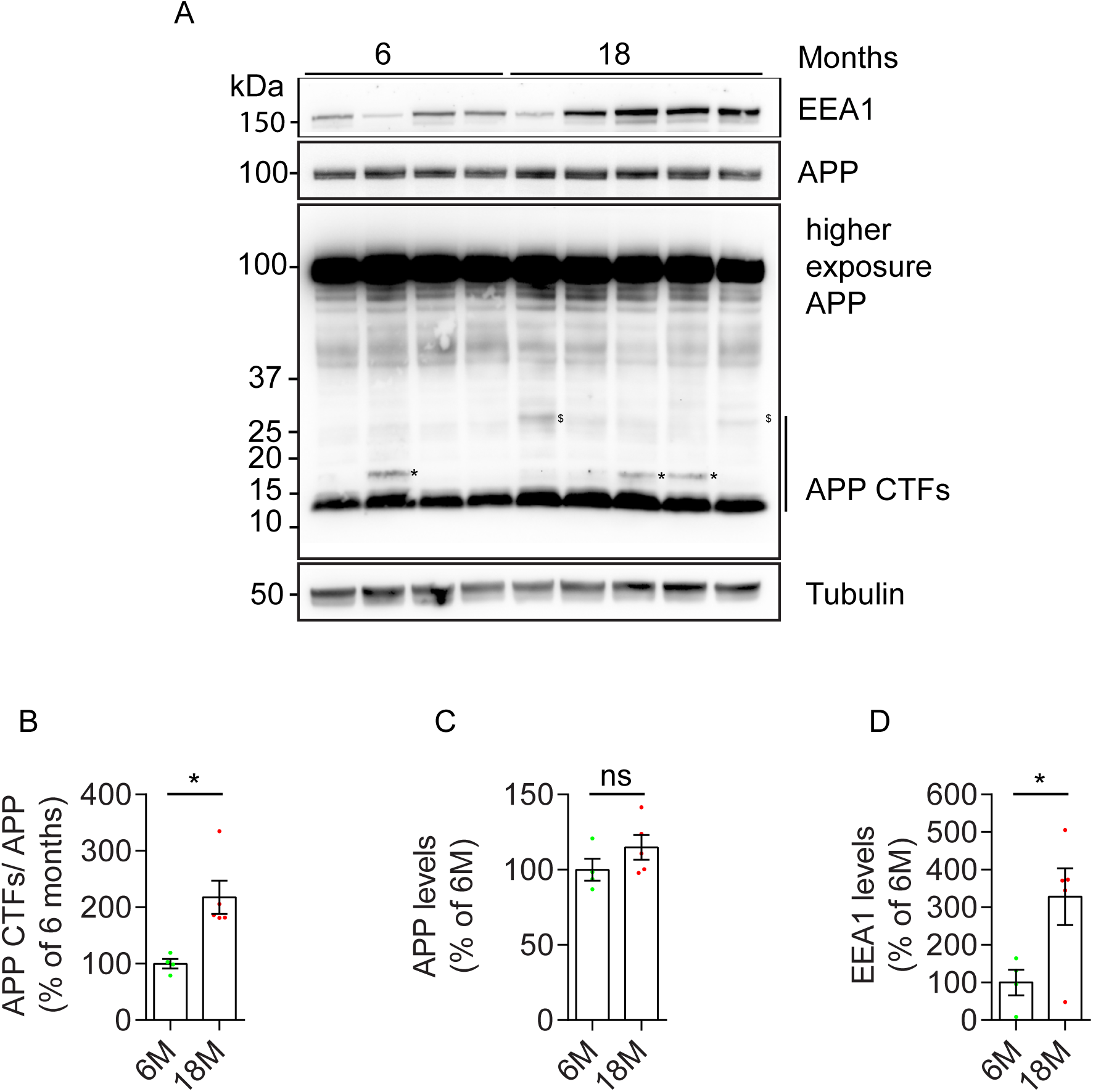
Aged brain evidences increased APP processing and early endosomes accumulation. A. Endogenous EEA1 levels (first panel), APP levels (second panel), APP-CTFs levels (third panel, higher exposure) and tubulin as loading control in adult (6 months old) and aged (18 months old) mice cortical brain analyzed by Western blot with anti-EEA1 antibody, anti-APP antibody (Y188) and anti-tubulin. Note: * indicate beta-longer CTF fragment (C99); $ indicate eta-CTF. B. Quantification of APP-CTFs levels normalized to APP and to percentage of 6 M (n = 4-5; \**P* = 0.0159 18 M vs. 6 M brain, Mann-Whitney test, mean ± SEM). C. Quantification of APP levels normalized to tubulin and to percentage of 6 M (n = 4-5; 18M APP level is not significantly different form 6 M *P* = 0.1905 18 M vs. 6M neurons, Mann-Whitney, mean ± SEM). D. Quantification of EEA1 levels normalized to tubulin and to percentage of 6 M (n = 4-5; 18M there is a tendency for EEA1 level to be higher at 18 M (*P* = 0.1111 18 M vs. 6M brains), Mann-Whitney, mean ± SEM).

Together these results indicate that, during aging, there is an up-regulation of endocytosis which correlates with increased processing of APP in the brain, recapitulating fundamental mechanistic findings in *in vitro* neurons aged.

### Aging-dependent synapse loss is in part due to Aβ production

Synapse dysfunction, more than neuronal death or dendritic shrinkage, has been observed in the aging brain. Synapse dysfunction is thought to account for the cognitive decline that the elderly develops, and it may precede AD. To determine if the increased Aβ production by aged neurons had a synaptic impact, we first assessed synapse decline in our neuronal model of aging, as previously reported both *in vitro* and *in vivo* (Nwabuisi-Heath et al., 2012; Petralia et al., 2014). As a proxy for a synapse, we imaged and quantified, the juxtaposition of the pre-synaptic marker vGlut1, a glutamate transporter, and of the post-synaptic marker PSD-95, a post-synaptic density scaffold that anchors glutamate receptors at synapses. Consistent with previous studies, we found that the density of synapses decreased significantly by 54% in aged neurons (Fig. 7A, 7B, 7D, 7E and 7M) (Papa et al., 1995; Nwabuisi-Heath et al., 2012). Given that the density of synapses declines in aged neurons that also show increased production of Aβ (see Fig. 1 and 2), we hypothesized that we could rescue the age-dependent reduction in synapse number by inhibiting γ-secretase-dependent Aβ production for 24 h with γ-secretase inhibitor (DAPT), which we previously showed to efficiently block APP processing (Almeida et al., 2005). Importantly, when we treated aged neurons with DAPT, the number of synapses increased significantly by 42%, partially rescuing the synapses lost due to neuronal aging (Fig. 7C, 7F, 7M). This result implicates Aβ42 production as a causal mechanism of synapse loss in aged neurons. Since the rescue was not complete, it indicates that other mechanisms of aging also contribute to synapse decline.

**Figure 7.**
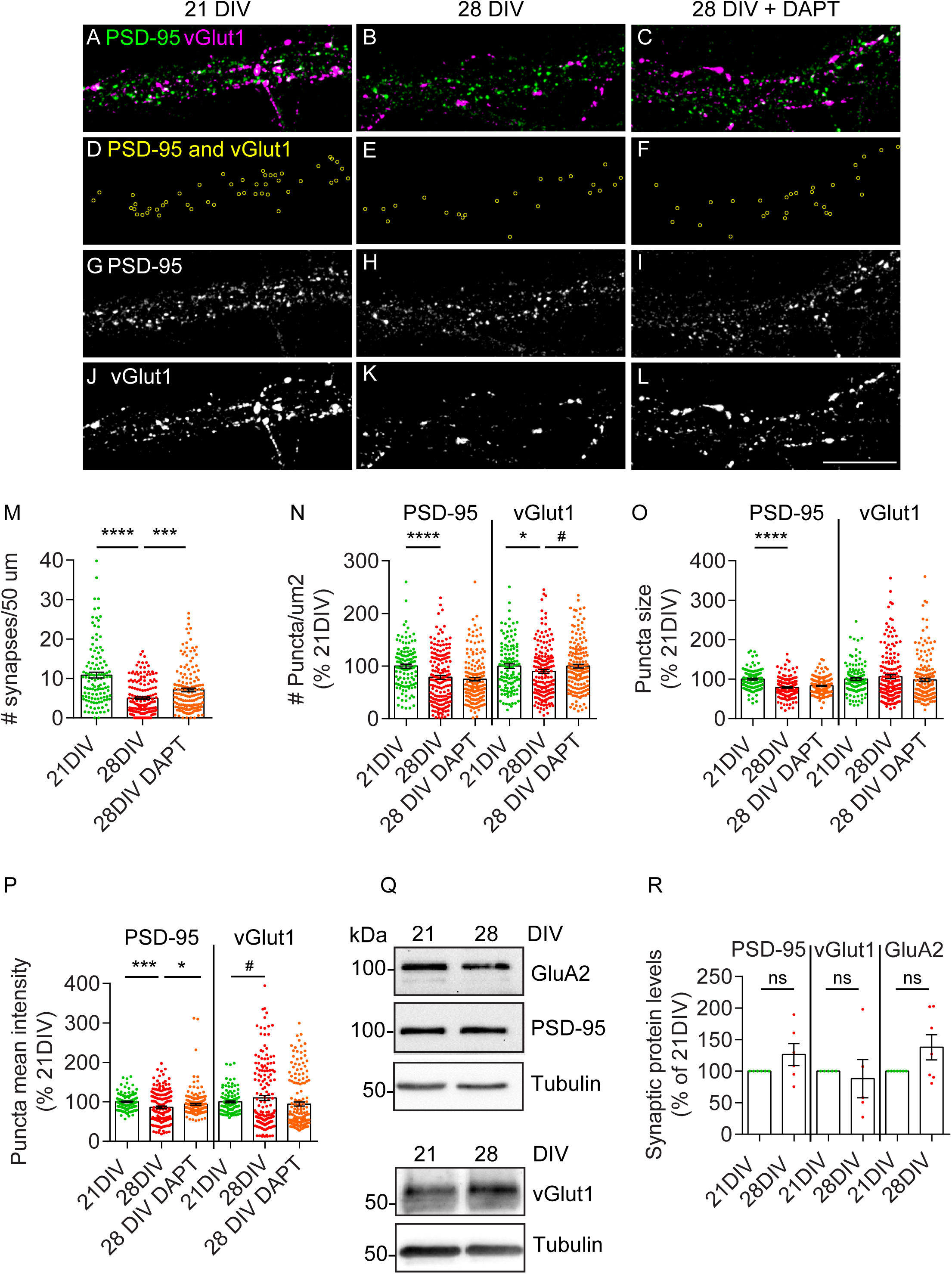
Aged neurons evidence synapse loss partially dependent on amyloid beta production. A-C. PSD-95 (green) and vGLut1 (magenta) detected by immunofluorescence in neurites of 21 DIV neurons (A), 28 DIV neurons (B) and 28 DIV neurons treated with DAPT (C), analyzed by epifluorescence microscopy and displayed after background subtraction with Fiji. Scale bar, 10 µm. D-F. Synapses (yellow rings) corresponding to juxtaposed PSD-95 and vGlut1 puncta’s in neurites of 21 DIV neurons (D), 28 DIV neurons (E) and 28 DIV neurons treated with DAPT (F) were generated automatically by ICY “Colocalizer” protocol. G-I. PSD-95 puncta in neurites of 21 DIV neurons (G), 28 DIV neurons (H) and 28 DIV neurons treated with DAPT (I) analyzed by epifluorescence microscopy and displayed after background subtraction with Fiji. Scale bar, 10 µm. J-L. vGlut1 puncta in neurites of 21 DIV neurons (J), 28 DIV neurons (K) and 28 DIV neurons treated with DAPT (L) analyzed by epifluorescence microscopy and displayed after background subtraction with Fiji. Scale bar, 10 µm. M. Quantification of the number of synapses per 50 µm of neurite of 21 DIV neurons (D), 28 DIV neurons (E) and 28 DIV neurons treated with DAPT (F) using ICY “Colocalizer” protocol (n=3; N_21DIV_=118 neurites; N_28DIV_= 161 neurites; N_28DIV DAPT_=155 neurites; \*\*\*\**P* < 0.0001 28 DIV vs. 21 DIV, \*\*\**P =* 0.0006 DAPT-treated vs. not treated 28 DIV, Mann Whitney test, mean ± SEM). N. Quantification of the number of PSD-95 and vGlut1 per 50 µm2 of neurites using ICY “Spot detector” plugin on 21DIV neurites, 28 DIV neurites and DAPT-treated 28DIV neurites. Results were normalized to percentage of 21 DIV (n=3; N_21DIV_=118 neurites; N_28DIV_=172 neurites; N_28DIV DAPT_=170 neurites; \*\*\*\**P*<0.0001 PSD-95 puncta density in 28 DIV vs. 21 DIV neurites, \**P=*0.0486 vGlut1 puncta density in 28 DIV vs. 21 DIV, ^#^*P=*0.0399 vGlut1 puncta density in DAPT-treated vs. not treated 28 DIV neurites, Mann Whitney test, mean ± SEM). O. Quantification of the size of PSD-95 and vGlut1 puncta using ICY “Spot detector” plugin on 21DIV neurites, 28 DIV neurites and DAPT-treated 28DIV neurites. Results were normalized to percentage of 21 DIV (n=3; N_21DIV_=118 neurites; N_28DIV_=172 neurites; N_28DIV DAPT_=170 neurites; \*\*\*\**P*<0.0001 PSD-95 size in 28 DIV vs. 21 DIV neurites, Mann Whitney test, mean ± SEM). P. Quantification of the mean intensity of PSD-95 and vGlut1 puncta using ICY “Spot detector” plugin on 21DIV neurites, 28 DIV neurites and DAPT-treated 28DIV neurites. Results were normalized to percentage of 21 DIV (n = 3; N_21DIV_ = 118 neurites; N_28DIV_ = 172 neurites; N_28DIV DAPT_ = 170 neurites; \*\*\**P*<0.0001 PSD-95 puncta mean intensity in 28 DIV vs. 21 DIV neurites, \**P =* 0.0283 PSD-95 puncta mean intensity in DAPT-treated 28 DIV vs. 28 DIV neurites, ^##^*P =* 0.0058 vGlut1 puncta mean intensity in 28 DIV vs. 21 DIV neurites, Mann Whitney test, mean ± SEM). Q. PSD-95, GluA2 and vGLut1 total levels in neurons at 21 DIV and 28 DIV analyzed by western-blot with anti-PSD-95 antibody, anti-GluA2 antibody and anti-vGlut1 antibody. Tubulin was immunoblotted as loading control. R. Quantification of PSD-95, GluA2 and vGLut1 levels normalized to tubulin and to percentage of 21 DIV (n_PSD-95_ = 6; n_vGlut1_ = 5; n_GluA2_ = 7; ns, not significant, Wilcoxon test, mean ± SEM).

To better understand why synapses were lost, we analyzed the density, size and intensity of each PSD-95 and vGlut1 puncta. We found that, for PSD-95, all three parameters were significantly reduced in aged neurons (Fig. 7G, 7H, 7N-7P). Surprisingly, only the intensity of PSD-95 puncta improved upon DAPT treatment (Fig. 7I, 7P). Further, vGlut1 density and intensity were slightly but significantly reduced in aged neurons (Fig. 7J, 7K, 7N, 7P). Interestingly, only the density of vGlut1 increased significantly upon DAPT treatment (Fig. 7L, 7N), indicating that Aβ production may impact both the PSD-95 positive post-synaptic compartment and the vGlut1 positive pre-synaptic compartment. Of note, the total levels of post-synaptic markers, PSD-95 and GluA2, the AMPA glutamate receptor subunit, as well as vGlut1 were not significantly altered in aged neurons (Fig. 7Q, R). Our data support that the defects observed refer to synapse loss and not to major neuronal degeneration (see Fig. 1).

Overall, from our data, we conclude that Aβ production may account at least in part for the detrimental effect of aging on synapses.

## Discussion

The accumulation of Aβ42 results from the imbalance between production and clearance. Aβ accumulation in the aging brain was until now explained by a decline in Aβ degradation with aging (Saido and Leissring, 2012); however, whether Aβ production increases with aging cannot be ruled out. Here, we observed that endogenous APP is increasingly processed with neuronal aging to generate Aβ. Importantly, we discovered that aged neurons endocytose more APP, facilitating APP processing required for Aβ generation in neurites. Furthermore, we also provide evidence that the normal age-related Aβ production is in part responsible for the loss of synapses by aged neurons, and thus may be initiating a pathological mechanism during aging which could trigger Alzheimer’s disease. Therefore, we propose that Aβ production is involved in neuronal aging.

### Intracellular Aβ42 and neuronal aging

Endogenous intracellular Aβ42 was found significantly increased in primary neurons aged in culture, which is consistent with previous observations of increased Aβ42 in conditioned media of aged neurons (Guix et al., 2012; Skovronsky et al., 1998; Kimura et al., 2005; Bertrand et al., 2011). These changes in Aβ42 accumulation with aging *in vitro* recapitulate the accumulation of intraneuronal Aβ42 (Blair et al., 2014; Baker-Nigh et al., 2015; Norvin et al., 2015; Kimura et al., 2005) and of amyloid plaques (Vlassenko et al., 2011; Petersen et al., 2016; Funato et al., 1998) in the normal aged brain.

Aβ42 increased predominantly in neurites, where most synapses occur. We had previously observed a similar accumulation of Aβ42 at synapses in AD-like transgenic neurons *in vitro* and *in vivo* (Takahashi et al., 2004, 2002; Tampellini and Gouras, 2010). This neurite-specific accumulation of Aβ42 could be explained by the increased presence of APP observed in aged neurites. APP delocalization to neurites could be due to a deficit in APP retrograde trafficking back to the cell body potentially caused by a possible malfunction of dynein with aging (Kimura et al., 2012, 2007).

While the degradation of Aβ has been reported to be decreased with aging, either due to reduced activity of Aβ degrading enzymes (Iwata et al., 2002) or decreased clearance by impaired glia phagocytosis (Solé-Domènech et al., 2016) and by reduced transport across the blood-brain barrier (Elahy et al., 2015), evidence supporting an increase in Aβ production with aging is scarce. We found that the intracellular accumulation of Aβ42 together with the augmented levels of APP CTFs both in *in vitro* aged neurons and in *in vivo* aged brain (Fig. 2 and Fig. 6) strongly indicate that APP processing originating Aβ42 is increased in aged neurons.

### Why does APP processing increase with aging?

Aging impact on APP processing is likely independent of alterations in the cellular levels of APP, which we found not to be significantly altered both *in vitro* and *in vivo*, supporting similar observations during normal brain aging (Flood et al., 1997; Gegelashvili et al., 1994). However, these results are not consensual since while some report APP increased with aging *in vitro* and *in vivo* (Sinha et al., 2016; Guix et al., 2012) other report APP decreased with aging *in vivo* (Kern et al., 2006). We hypothesize that these differences may occur due to the different brain regions analyzed or to the different conditions of primary neurons culture. Overall, one may conclude that the differences in APP levels, if exist, are not major and thus are not sufficient to account for the increase in APP processing with aging.

Increased APP processing may in part result from an aging-dependent increase in APP secretases activity (Guix et al., 2012; Fukumoto et al., 2004). We and others found that nicastrin levels are not altered by *in vitro* neuronal aging (Guix et al., 2012), although have also been described to be reduced *in vivo* (Placanica et al., 2009). Less is known about the trafficking of γ-secretase. A report indicates that the subcellular localization of nicastrin determines the site of assembly of γ-secretase (Morais et al., 2008) but whether it changes with aging is not known. Interestingly, nitrosative stress affects γ-secretase function during neuronal aging *in vitro* (Guix et al., 2012) which *in vivo* may contribute to increase Aβ42 (Placanica et al., 2009). The increase in BACE1 activity is not due to an increase in the level of BACE1 in the aged brain (Fukumoto et al., 2004), which we also did not see altered in our aged neurons. Instead, an increase in the access of APP to BACE1, due to altered APP trafficking, could underlie the increase in processing.

Indeed we discovered that APP endocytosis is significantly increased in aged neurons, which could explain the increase in APP processing, since endocytosis has been shown to be required for APP encounter with its secretases and for Aβ production (Guimas Almeida et al., 2018; Cirrito et al., 2008; Zou et al., 2007; Rajendran et al., 2008; Grbovic et al., 2003). Indeed, we show that in aged neurons, APP localization to early endosomes increased. Our findings, together with several other reports, support that early endosomes are the main site for the encounter of APP with its secretases and also for APP processing likely occurring during endosomal maturation into late-endosomes, where Aβ is known to accumulate (Almeida et al., 2006; Yuyama and Yanagisawa, 2009; Sannerud et al., 2016; Willén et al., 2017; Edgar et al., 2015; Takahashi et al., 2004, 2002; Morel et al., 2013; Rajendran et al., 2006; Vetrivel and Thinakaran, 2006). Thus, we identify endocytosis as a new mechanism whereby APP processing might increase with aging.

### How does APP endocytosis increase with aging?

While no previous study observed directly that APP endocytosis increases with neuronal aging, indirect evidence such as the presence of enlarged endosomes early in sporadic AD patient brains support that endocytosis is increased with aging (Cataldo et al., 1997, 2000b). Moreover, an increase in endocytic protein levels with aging has been reported (Blanpied et al., 2003; Alsaqati et al., 2017; Kimura et al., 2012). Interestingly, PICALM, a component of the clathrin-mediated endocytosis machinery involved in APP endocytosis (Xiao et al., 2012), which has been genetically associated with AD (Harold et al., 2009; Carmona et al., 2018), was reported to be increased with aging *in vivo* (Alsaqati et al., 2017).

Contrary to the generalized increase in endocytosis with aging is the fact that we found transferrin endocytosis reduced in aging neurons, which could be due to a reduced rate of transferrin exit from endocytic clathrin-coated pits in aged neurons (Blanpied et al., 2003). Alternatively, the reduced transferrin endocytosis might be due to a specific reduction in the expression of transferrin receptor in the aged hippocampus (Lu et al., 2017).

Endocytosis of APP, like of transferrin receptor, is mostly clathrin-mediated (Koo and Squazzo, 1994; McMahon and Boucrot, 2011). To identify the mechanisms which underlie the up-regulation of APP endocytosis and the down-regulation of transferrin endocytosis more research is necessary. These apparently contradictory changes indicate that specific mechanisms of APP endocytosis may arise during neuronal aging and ignite a new research field.

### Early endosome up-regulation with neuronal aging

The endocytosis of APP allows it to enter early endosomes that harbor Rab5 and its effector EEA1. In aged neurons, the up-regulation of APP endocytosis was accompanied by an increase in early endosomes marked by EEA1. Early endosomes of aged neurons were bigger and brighter, especially in neurites. Also, in the aged brain, we found that the early endosome marker EEA1 was elevated. This up-regulation of early endosomes had been previously observed in *in vitro* aged neurons and in sporadic AD (Blanpied et al., 2003; Cataldo et al., 2000a). Early endosomes form by fusion of endocytic vesicles formed upon endocytosis. Thus, the up-regulation of early endosomes could be due to the increase in APP endocytosis. Still, we cannot exclude the contribution of a deficit in early endosome maturation and in lysosomal degradation with neuronal aging.

On a different perspective, the recruitment of EEA1 to early endosomes could be due to the increased activation of Rab5, an early endosome specific small GTPase and a major regulator of endocytosis (Rubino et al., 2000; Rybin et al., 1996; Christoforidis et al., 1999; Bucci et al., 1992; Alsaqati et al., 2017). Indeed, Rab5 itself has been found up-regulated with aging (Ginsberg et al., 2011; Neefjes and van der Kant, 2014). In addition, an impairment of EEA1 proteostasis may underlie its overall increased levels in aged neurons and in the aged brain. Research is needed to determine the mechanisms of EEA1 proteostasis normally and with aging.

Overall, our findings indicate that early endosome enlargement results from increased endocytic uptake.

### Synapse dysfunction in aging and Alzheimer’s

Although the accumulation of Aβ is concomitant with synapse dysfunction in the aging brain, causality has not been established. Here, we found that inhibition of Aβ production increases the number of synapses in aged neurons. However, the rescue was only partial, indicating that other mechanisms could contribute to synapse loss during aging (Yankner et al., 2008).

Interestingly, the fact that aged neurons show Aβ dependent synapse loss after 28 DIV while previously we had demonstrated that Aβ overproduction in familial AD neurons, driven by the Swedish mutation in APP, present synapse loss after only 19 DIV (Almeida et al., 2005) indicates that, as expected, the production of Aβ by aged neurons is inferior than by familial AD neurons driving a slower synapse loss with aging. Moreover, our data indicating that increased Aβ production with aging is impacting both the PSD-95 positive post-synaptic compartment and the vGlut1 positive pre-synaptic compartment is in agreement with our previous data in fAD neurons (Almeida et al., 2005; Takahashi et al., 2004).

Overall, our findings indicate that the up-regulation of APP endocytosis is a cell autonomous mechanism by which neurons contribute to brain aging. Our next step will be to device new technology that allow for measuring endocytosis in neurons *in vivo*. It will be interesting to determine the impact of aging on other brain cells, such as astrocytes and microglia. We plan in our future studies to incorporate several brain cell-types in more complex culture models to study brain aging mechanisms.

Our work highlights the involvement of APP endocytosis as a key link between aging and AD. The increased processing of APP and Aβ42 accumulation points to an impairment in endocytic trafficking towards early endosomes as neurons get older. Also, we found that the detrimental effect of aging on synapses account, at least in part, to Aβ production. Hence, the identification of the mechanisms underlying aging-synaptic decline is urgent to reverse synaptic dysfunction and thus prevent cognitive decline in the face of aging, delaying AD.

## Materials and Methods

### Cell culture

Primary neuronal cultures were prepared as previously reported (Almeida et al., 2005) from cortices of embryonic day 16 (E16) wild-type females and males BALB/c mice (Instituto Gulbenkian Ciência and CEDOC). All animal procedures were performed according to EU recommendations and approved by: Instituto Gulbenkian de Ciência Animal Care and Ethical Committee; the NMS-UNL ethical committee (07/2013/CEFCM) and the national DGAV (0421/000/000/2013). Briefly, E16 brain tissue was dissociated by trypsinization and trituration in DMEM with 10% fetal bovine serum (Heat-Inactivated FBS, Life Technologies). Dissociated neurons plated in DMEM with 10% FBS on poly-D-lysine (Sigma-Aldrich)-coated 6-well plates (1 × 10^6^ cells/cm2) and glass coverslips (5 × 10^4^ cells/cm2). After 3-16 h media was substituted for Neurobasal medium supplemented with B27, GlutaMAX and penicillin/streptomycin (all from Life Technologies) at 37 °C in 5 % CO2. Cells were maintained up to 28 days in vitro (DIV) without changing or adding new media.

When indicated, γ-secretase was inhibited by 24 h of treatment with 250 nM DAPT (Calbiochem) and DMSO (solvent) was used as control.

### Antibodies

The following primary antibodies were used: Anti-Ankyrin-G pAb (P-20, Santa Cruz, cat sc-31778, 1:100); anti-APP mAb (22C11, Millipore, cat MAB348, 1:100); anti-APP (Y188, GeneTex, cat GTX61201, 1:200; 1:1,000); anti-Aβ42 mAb (12F4, Millipore, cat 05-831-l, 1:50); anti-BACE1 pAb (Thermo Scientific, cat PA1-757, 1:850); anti-EEA1 pAb (N-19, Abcam, cat sc-6415, 1:50); anti-MAP2 mAb (Sigma, M4403, 1:500); anti-nicastrin pAb (Thermo Scientific, cat PA1-758, 1:500); anti-tubulin mAb (Tu-20, Millipore, cat MAB1637, 1:10,000); anti-LAMP1 (CD107a, BD Pharmingen, cat 553792; 1:200); anti-Rab5 (Sicgen, cat AB1024-200, 1:200); anti-GAPDH (Ambion, cat AM4300, 1:1000); anti-PSD-95 (Merck Millipore, cat 04-1066, 1:200; 1:1,000); anti-vGlut (Merck Millipore, cat MAB5502, 1:200; 1:1,000); anti-GluR2 (Merck Millipore, cat MABN71, 1:1000). The secondary antibodies used were conjugated to Alexa-488, −555 and −647 (Molecular Probes) or to HRP (Bio-rad).

### Immunofluorescence labelling

Immunofluorescence was performed as previously (Almeida et al., 2005; Ubelmann et al., 2017b). Briefly, cultured primary neurons were fixed at 14, 21 and 28 days *in* vitro (DIV) with 4% paraformaldehyde/4% sucrose in PBS for 20 min, permeabilized with 0.1% saponin in PBS for 1 h and blocked in 2% FBS/1% BSA/0.1% saponin in PBS for 1 h at room temperature (RT) before antibody incubation using standard procedure. For PSD-95 and vGlut1 immunolabelling, permeabilization was performed with 0.3 % Triton-X in PBS for 5 min at RT. For APP surface labelling, cells were incubated with primary and secondary antibodies prior to permeabilization and immunolabelling (Ubelmann et al., 2017a). Coverslips were then mounted using Fluoromount-G (Southern Biotechnology, Birmingham, AL).

### Image acquisition

Epifluorescence microscopy was carried out on an upright microscope Z2 (Carl Zeiss) equipped a 60× NA-1.4 oil immersion objective and an AxioCam MRm CCD camera (Carl Zeiss) or on an upright microscope DMRA2 (Leica) equipped with a 100× NA-1.4 oil immersion objective and a CoolSnap HQ camera (Photometrics). Confocal microscopy was performed with LSM710 (Zeiss) or with a Revolution xD (Andor) spinning-disk system coupled to an Eclipse Ti-E microscope (Nikon). For direct comparison, samples were imaged in parallel and using identical acquisition parameters.

### Immunoblotting

Cell lysates were prepared using modified RIPA buffer (50 mM Tris– HCl pH 7.4, 1 % NP-40, 0.25 % sodium deoxycholate, 150 mM NaCl, 1 mM EGTA, 0.1 % SDS, with PIC). Proteins separated by 7.5, 10 or 15% Tris-glycine SDS–PAGE or 4-12% Tris-Glycine SDS-PAGE (Invitrogen) were transferred to nitrocellulose membranes and processed for immunoblotting using ECL Prime kit (GE Healthcare). Images of immunoblots were captured using ChemiDoc (Bio-Rad) within the linear range and quantified by densitometry using the “Analyse gels” function in ImageJ.

### Trafficking assays

For bulk endocytosis of biotinylated APP (Fig. 4) biotinylation of surface APP was performed as previously described (Almeida et al., 2006). Briefly, neurons at 21 DIV and 28 DIV were incubated on ice with 0.5 mg/ml Sulfo-NHS-LC-Biotin (Pierce) in PBS for 30 min. Free biotin was quenched with ice-cold 0.5 % BSA in PBS. Biotinylated proteins were chased for 10 min and 30 min at 37 °C to allow for detecting endocytosis. For detection of surface biotin-APP, cells were rinsed, and lysates prepared in RIPA buffer. For detection of endocytosed biotin-APP, surface biotin was stripped by treating cells with GSH (50 mM) in stripping buffer (75 mM NaCl, 10 mM EGTA, 1% BSA, pH 7.8 - 8) for 15 min on ice before lysates were prepared. Biotinylated proteins were immunoprecipitated with NeutrAvidin agarose beads (Pierce) overnight at 4°C and, after washing, separated by SDS-PAGE. Quantitative immunoblotting was performed using anti-APP (Y188) antibody in biotinylated proteins and total proteins.

For APP endocytosis (Fig. 4), a 10 min pulse with a monoclonal mouse antibody against the extracellular N-terminus of surface APP (22C11) was performed as previously described (Ubelmann et al., 2017a). After a 10 min pulse at 37°C in complete medium with 10 mM HEPES, cells were fixed and immunolabelled with a secondary antibody anti-mouse and mounted or co-labelled for Rab5 (Fig. S2).

For transferrin endocytosis (Fig. 5), a 10 min pulse with transferrin was performed as previously described (Almeida 2006). Briefly, neurons were incubated with 10 µg/ml of transferrin labelled with Alexa647 (A647-Tf; LifeTechnologies, cat. T23366) in complete medium with 10 mM HEPES for 10 min (pulse) at 37°C. After, cells were fixed, permeabilized and labelled with Phalloidin to probe F-actin, washed and mounted.

### Quantitative analyses

Image analyses were carried out using ImageJ (imagej.nih.gov/ij), Fiji (fiji.sc) or ICY (icy.bioimageanalysis.org).

For the quantification of the number of auto-fluorescent granules positive for LAMP1 (Fig. 1K) and auto-fluorescent granules (Fig. S1) per area of dendrite (density), dendritic segments were outlined with ICY, LAMP1 and auto-fluorescent granules were segmented and counted automatically using the ICY “Spot detector” plugin, and the number of colocalizations (lysosomal lipofuscin) was obtained using the ICY “colocalization studio” plugin.

For the subcellular quantification of intracellular Aβ42 levels (Fig. 1R) and of APP levels (Fig. 3C) the mean fluorescence of Aβ42/APP in the cell body and in neurites was measured using ImageJ. The cell body and a region of background was outlined using “polygon selection”. For selecting a region of neurites, a square (500 x 500) was centered on each primary dendrite. The mean fluorescence of Aβ42/APP in each region was quantified with “Measure” function. The mean fluorescence per region was calculated as percentage of the indicated control, upon background fluorescence subtraction.

For the quantification of APP levels in axons vs dendrites (Fig. 3J, K), two subcellular regions of interest (ROI), axon (AnkG positive) and dendrite (AnkG negative) were outlined using ImageJ “polygon selection”. The mean fluorescence of APP in each ROI was quantified as above. For the quantification of APP polarization, APP mean fluorescence in the axon ROI was divided by the APP mean fluorescence in the dendrite ROI (APP axon/dendrite ratio). APP mean fluorescence in axons vs dendrites was calculated as above.

For the quantification of APP colocalization with EEA1 density per area (Fig. 3P), of endocytosed APP (22C11) colocalization with Rab5 percentage (Fig.S2), or PSD-95 colocalization with vGlut1 (synapse) density per length of neurite (Fig. 7M), the number of colocalizing objects was obtained using ICY “Colocalizer” protocol. The area or length (Feret’s diameter) of the neurite ROI was obtained using ICY ROI export.

For the quantification of puncta density per area, intensity and size of endocytosed APP (22C11), EEA1, PSD-95, vGlut (Fig. 4M, 4N, 4O, 5G, 5H, 5I, 7N, 7O, 7P) and for density of endocytosed transferrin (Fig. 5Q), the ICY “spot detector” was used.

### Brain homogenates preparation

Adult (6 months) and aged (18 months) mice brains were solubilized using modified RIPA buffer. After 15 min on ice with RIPA buffer, brains were sonicated and centrifuged at 4 °C for 30 min, 13.3 rpm. Equivalent amounts of protein (40 µg), as determined by the BCA protein assay kit (Thermofisher), were mixed with sample buffer, heated at 95°C for 5min, vortexed, and run on 4-12% Tris-Glycine SDS-PAGE (Invitrogen). Electrophoretic transfer and immunoblot was performed as described in immunoblotting.

### Statistics

GraphPad Prism 6 software was used for graphs generation of individual replicates with mean ± SEM and for statistical analysis of at least three independent experiments as indicated in figure legends. Sample size was determined based on pilot studies. Data was tested with D’Agostino-Pearson omnibus normality test. For non-parametric and paired data, the Wilcoxon t test was applied; for non-parametric and unpaired data the Mann-Whitney (shift in median) or Kolmogorov-Smirnov (shift in distribution) test was applied; for non-parametric and using multiple comparisons statistical analysis of data the one-way ANOVA on ranks with *post hoc* Dunn’s testing was applied, as specified in figure legends.

## Acknowledgements

We thank for the gift of antibodies to Dr. Stephanie Miserey-Lenkei (Institute Curie). We thank Dr. F. Ubelmann, Dr. R. M. Oliveira and L. Salavessa for helpful discussions. We thank M. Guimarães and D. Rodrigues for their technical assistance with synaptic markers immunofluorescence assays in neurons and L. Almeída for an excel macro. We thank the lab members for helpful discussions and for critical reading of the manuscript. We thank Dr. H. V. Miranda (CEDOC), Dr. A. Gontijo (CEDOC) and Dr. E. Gomes (IMM) for critical reading of the manuscript.

We thank Dr. M. Rebelo (IGC Animal Facility) and Dr. S. Marques (CEDOC Animal Facility). We thank Dr. N. Pimpão (IGC) and Dr. T. Pereira (CEDOC) for their expertise in microscopy. This work has been supported by: Marie Curie Integration Grant (PCIG-GA-2012-334366-trafficinAD; Marie Curie Actions, EC); iNOVA4Health—UID/Multi/04462/2013, a program financially supported by Fundação para a Ciência e Tecnologia (FCT)/ Ministério da Educação e Ciência, through national funds and co-funded by FEDER under the PT2020 Partnership Agreement); Maratona da Saúde 2016; Investigator FCT (IF/00998/2012, FCT). TB has been the recipient of an FCT doctoral fellowship (SFRH/BD/131513/2017). AT was the recipient of an FCT doctoral fellowship (SFRH/BD/52473/2014).

## Author contributions

Conceptualization, Project management, Funding acquisition, and supervision: C. Almeida; Formal analysis: C. Almeida, T. Burrinha (lead), R. Gomes; Funding acquisition: C. Almeida; Investigation: T. Burrinha (lead), C. Almeida, R. Gomes, A. P. Terrasso; Methodology: C. Almeida, T. Burrinha. Visualization and writing: C. Almeida, T. Burrinha.

