## supplemental figures for "Neuronal aging potentiates beta-amyloid generation via amyloid precursor protein endocytosis"

Figure S1

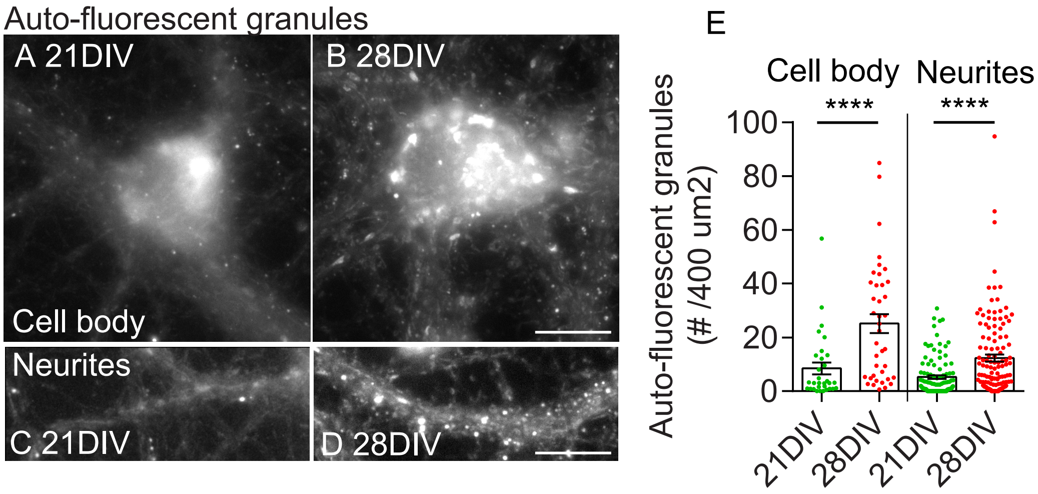

Auto-fluorescent granules
A 21DIV
B 28DIV
Cell body
Neurites
C 21DIV
D 28DIV
E
Cell body
Neurites
****
****
Auto-fluorescent granules
(# /400 um2)
100
80
60
40
20
0
21DIV
28DIV
21DIV
28DIV

Figure S2

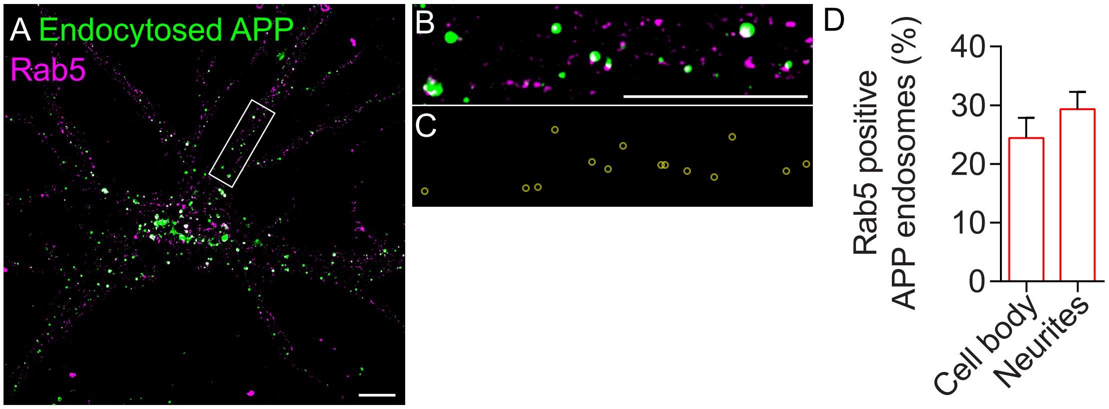
A Endocytosed APP
Rab5
B
C
D
Rab5 positive
APP endosomes (%)
40
30
20
10
0
Cell body
Neurites

**Figure S1**. Aged neurons evidence canonical signs of aging, such as the accumulation of auto-fluorescent granules.

A-D. Auto-fluorescent aging granules, or lipofuscin, in cell bodies of 21DIV (A) and 28DIV (B) neurons and neurites of 21DIV (C) and 28DIV (D) neurons analyzed by epifluorescence microscopy. Scale bars, 10 µm.

E. Quantification of the number of auto-fluorescent granules per area (400 µm2) in cell bodies and neurites (n=3-4, $N_{cellbody}$ = 31-39, $N_{neurites}$= 60-129, ****$P_{cellbody}$ < 0.0001 28 DIV vs. 21 DIV cell body, ****$P_{neurites}$ < 0.0001 28 DIV vs. 21 DIV neurites, Mann-Whitney test, mean ± SEM).

**Figure S2**. Endocytosed APP is present in Rab5 positive endosomes at 28DIV neurons.

A-C. Endocytosed APP (10 min 22C11; green) and Rab5 (magenta) in neurons was assessed by immunofluorescence at 28 DIV (A), with anti-Rab5 (Rab5; magenta) and analyzed by epifluorescence microscopy. Images are displayed upon background subtraction. The white rectangle indicates the magnified neurite (B). Rab5-positive APP endosomes (yellow rings) on the magnified neurite were generated automatically by ICY “Colocalizer” protocol (C). Scale bar, 10 µm.

D. Quantification of the number of Rab5-positive APP endosomes in the cell body and neurites in percentage of endocytosed APP puncta (n=2, $N_{cellbody}$ = 12, $N_{neurites}$= 36).
